# Prime editing outperforms homology-directed repair as a tool for CRISPR-mediated variant knock-in in zebrafish

**DOI:** 10.1101/2025.02.05.636566

**Authors:** Michiel Vanhooydonck, Elyne De Neef, Hanna De Saffel, Annekatrien Boel, Andy Willaert, Bert Callewaert, Kathleen BM Claes

**Affiliations:** Center for Medical Genetics, Ghent University Hospital, 9000 Ghent, Belgium; Department of Biomolecular Medicine, Ghent University, 9000 Ghent, Belgium; CRIG (Cancer Research Institute Ghent), 9000 Ghent, Belgium; Ghent-Fertility and Stem Cell Team (G-FaST), Department for Reproductive Medicine, Ghent University, 9000 Ghent, Belgium

**Author notes:** Corresponding Authors: Kathleen BM Claes, Bert Callewaert. Michiel Vanhooydonck and Elyne De Neef contributed equally to this work. Andy Willaert, Bert Callewaert and Kathleen BM Claes contributed equally as senior authors to this work.

## Abstract

Zebrafish serve as a valuable model organism for studying human genetic diseases. While generating knockout lines is relatively straightforward, introducing precise disease-specific genetic variants by knock-in (KI) remains challenging. KI lines, however, enable more accurate studies of molecular and physiological consequences of genetic diseases. Their generation is often hampered by low editing efficiencies (EE) and potential off-target effects. In this study, we optimized conventional CRISPR/Cas9-mediated homology-directed repair (HDR) strategies for precise KI of genetic variants in zebrafish and compared their efficacy with prime editing (PE), a recently developed technique that is not yet commonly used. Using next-generation sequencing (NGS), we determined KI EE by HDR for six unique base-pair substitutions in three different zebrafish genes. We assessed the effect of 1) varying Cas9 amounts, 2) HDR templates with chemical modifications to improve integration efficiency, 3) different micro-injection procedures, and 4) synonymous guide-blocking variants in the protospacer sequence. Increasing Cas9 amounts augmented KI EE, with optimal injected amounts of Cas9 between 200 and 800 pg. The use of Alt-R™ HDR templates (IDT) further increased KI EE, while guide-blocking modifications did not. Injecting components directly into the cell was not superior to injections into the yolk. PE, however, increased EE up to fourfold and expanded the F0 founder pool for four targets when compared to conventional HDR editing, with fewer off-target effects. Therefore, PE is a very promising methodology for improving the creation of precise genomic edits in zebrafish, facilitating the modeling of human diseases.

## INTRODUCTION

Over the past decade, zebrafish (*Danio rerio*) have emerged as a valuable model for studying gene functions. Zebrafish possess at least one ortholog for 70% of all human genes and 80% of known human disease-causing proteins. Their high fecundity, external and rapid development, short reproductive cycle, and *ex utero* development contribute to their popularity in disease modeling^1–3^. Knockout (KO) models, where the function of a gene of interest is disrupted by introducing a truncating variant, are valuable and widely used for studying gene function. However, the need for precise knock-in (KI) models is increasing to overcome the intrinsic limitations of KO models. Even when aiming for loss of function (LOF) of a gene, using a KI approach may have advantages. Premature termination codons, which usually result in mutant mRNA degradation, can potentially trigger transcriptional adaptation of compensatory genes and pathways, (partially) rescuing the function of the targeted gene and effectively compensating for its loss^4–6^. Using KI strategies to mutate a key residue or create an in-frame deletion avoids nonsense-mediated decay and transcriptional adaptation. Furthermore, precise KI models are required to study the effects of single nucleotide variants resulting in gain-of-function, dominant negative effects, or to circumvent lethality, potentially associated with complete KO^7^. Finally, interest is growing in using zebrafish as a model to evaluate variants of unknown significance or variants that may affect splicing in disease. Therefore, mimicking human diseases as closely as possible by knocking-in specific variants in an animal model is essential for translational research approaches.

Conventional CRISPR/Cas9-based methods to generate zebrafish KI models use a crRNA:tracrRNA complex to guide Cas9 to a specific genomic location, where it introduces a double-strand break (DSB). In the presence of a homology-directed repair (HDR) template containing the variant of interest, the HDR pathway can be activated, allowing the variant to be introduced into the genomic sequence^8^. Various types of HDR templates have been utilized, including circular and linear double- or single-stranded templates^9,10^.

For KI generation, both double-stranded and single-stranded templates are commonly used. However, single-stranded HDR templates are preferred due to their higher efficiency^11–13^, lower susceptibility to nonspecific integration^14^, and straightforward design and production^15^. The length of the HDR arms impacts editing efficiency (EE) in zebrafish^16^. Our group previously optimized HDR template length, identifying the optimal length to be around 120 nucleotides^15^. However, for other loci, different HDR template lengths have been shown to be optimal^17,18^. Finally, chemical modifications of the HDR template ends can improve EE. Prykhozhij *et al.* (2018) demonstrated that phosphorothioate linkages at oligo ends enhance KI efficiency by reducing the impact of exonuclease activity on the template^19^.

Despite recent advancements in zebrafish genome editing methods, various parameters have not been systematically optimized or validated, with methodologies often relying on local expertise and the preferences of reporting laboratories. These parameters include the amount of Cas9 protein used, chemical modification of the HDR template ends, integration of silent guide-blocking mutations into the HDR template, and the injection of ribonucleoproteins (RNP) into the cell versus the yolk. Nevertheless, the process of KI generation in zebrafish typically remains inefficient^16^, compelling researchers to optimize innovative methods in zebrafish research. One such method is prime editing (PE), which does not require the generation of potentially harmful DSBs^20^. The prime editing guide RNA (pegRNA) directs the PE complex, consisting of Cas9 nickase and reverse transcriptase, to the target locus. The PE complex creates a single-stranded DNA nick. The primer binding site (PBS) hybridizes with the target DNA and serves as primer to initiate reverse transcription, during which the reverse transcriptase template (RTT) containing the desired edit is incorporated as DNA into the target region. Multiple DNA hybridization events between the pegRNA and the genomic target enhance the specificity to incorporate the RTT template. This technique further bypasses error-prone repair pathways like non-homologous end joining (NHEJ), resulting in lower off-target editing activity^20,21^.

In this study, we examined various parameters that could enhance the efficiency of traditional homology-directed repair (HDR) in zebrafish, which serves as a model system for gene editing. To this end, we knocked-in six unique missense variants in three different zebrafish genes. Additionally, we explored PE as a potentially more efficacious alternative.

## RESULTS

### Selection of knock-in targets in the zebrafish genome

In this study, we optimized KI strategies for six unique variants in three zebrafish genes associated with Mendelian disorders in humans: *col1a2*, *atp6v1e1b* and *brca2,* located on chromosomes 19, 4 and 15 and associated with osteogenesis imperfecta/Ehlers-Danlos syndrome, autosomal recessive cutis laxa type II C, and breast and ovarian cancer respectively. In our previous work, we characterized KO zebrafish models for the *col1a2*^22^, *atp6v1e1b*^7^ and *brca2*^23,24^ genes. We have now developed zebrafish KI models for two known benign human *COL1A2* variants: c.948C>T/p.(Gly316=) and 1446A>C/p.(Pro482=), by modifying the homologous zebrafish residues, respectively c.918A>T/p.(Gly306=) and c.1416A>C/p.(Pro472=). Additionally, we introduced the *col1a2* c.2093G>A/p.(Arg698Gln) variant, which is homologous to the human variant of uncertain significance (VUS), c.2123G>A/p.(Arg708Gln). Furthermore, we created zebrafish models for the *atp6v1e1b* c.634_636CGA>TGG/p.(Arg212Trp) and c.383T>C/p.(Leu128Pro), which cause the same amino acid changes as the human pathogenic variants c.634C>T/p.(Arg212Trp) and c.383T>C/p.(Leu128Pro), respectively^25^. In the *brca2* gene, we knocked-in the c.6803G>A/p.(Asp2268Asn) variant, which results in the same amino acid change as the human *BRCA2* variant c.8167G>A/p.(Asp2723Asn), a variant reported as damaging and located in a conserved residue of the DNA binding domain^26^. These KI zebrafish models may be utilized to investigate the phenotype and impact on gene function, or for functional testing of VUS in the future.

### Optimal Cas9 amount for homology-directed repair in zebrafish embryos is target-dependent

To determine the optimal Cas9 amount for each target, we tested four different amounts (100, 200, 400 or 800 pg) in combination with Alt-R™-modified HDR templates. These complexes were injected in triplicate into the yolk of single-cell zebrafish embryos. The highest EE, ranging from 0.8% to 5.0%, were achieved with 200 pg Cas9 for *col1a2* c.918A>T, 400 pg Cas9 for *col1a2* c.1416A>C and *col1a2* c.2093G>A, and 800 pg Cas9 for *atp6v1e1b* c.634_636CGA>TGG, *atp6v1e1b* c.383T>C and *brca2* c.6803G>A. Indel frequency (IF) increased with higher Cas9 amounts, with the highest IF observed at 800 pg Cas9, except for *col1a2* c.1416A>C and *col1a2* c.918A>T, which showed the highest IF at 400 pg and 200 pg, respectively (**Figure 1, Supplementary Table 1**).

**Figure 1.**
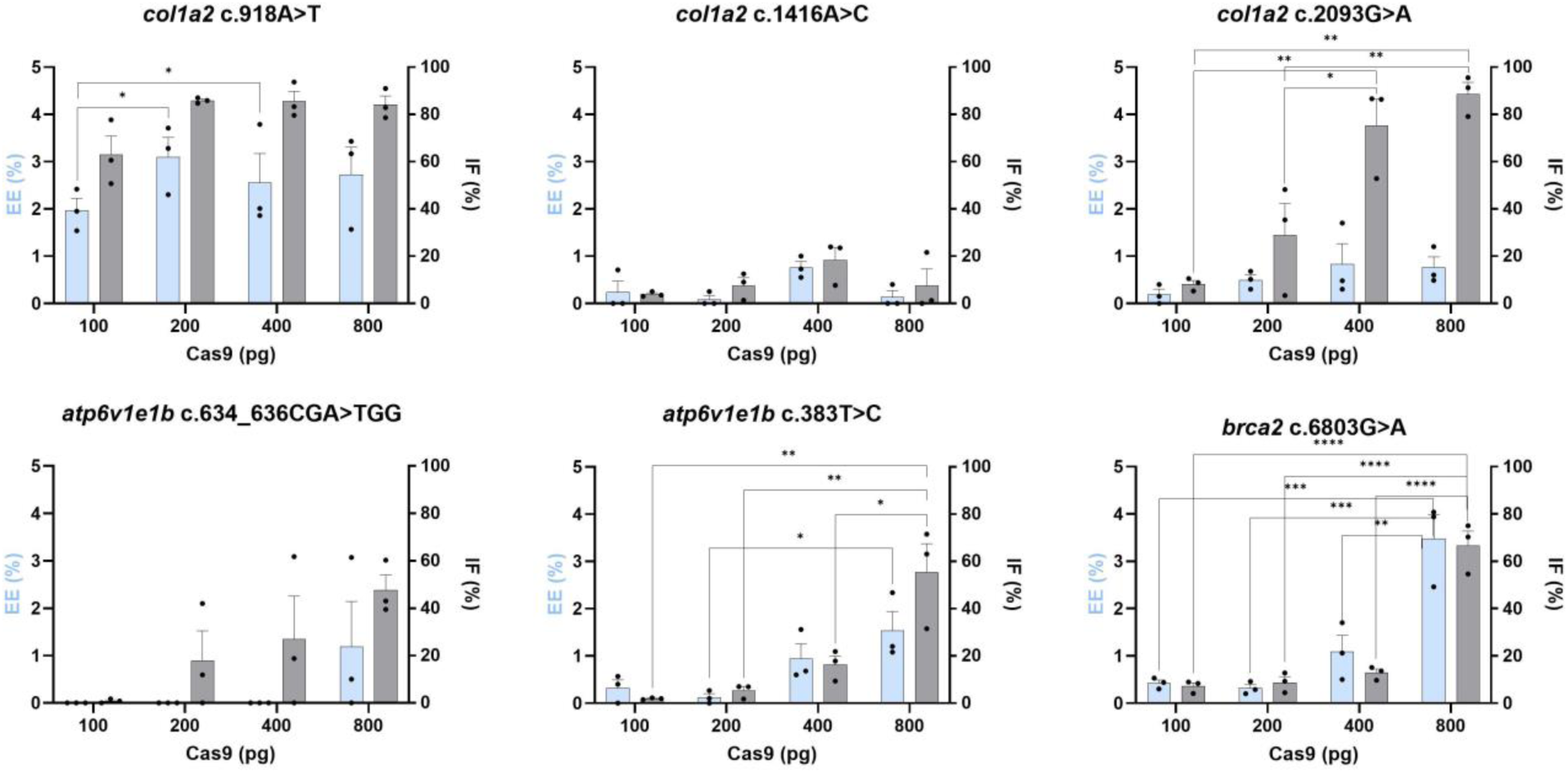
Editing efficiencies (EE - blue) and indel frequencies (IF - grey) obtained by microinjection of Alt-R™ HDR components with 100, 200, 400 or 800pg Cas9 into the yolk of 1-cell stage zebrafish embryos. Bars and error bars represent the mean + SEM of 3 biologically independent replicates (n=3), where each replicate represents a pool of 20-30 injected embryos. Tukey’s Multiple comparisons test: *p < 0.05; **p < 0.01; ***p < 0.001.

### Similar editing efficiencies achieved with yolk and cell injection

Previous studies have suggested that the highest EE can be achieved through direct microinjection into the cell of one-cell stage zebrafish embryos^16^. However, this method is more laborious, time-consuming, and technically challenging compared to injection into the yolk, as each embryo must be carefully aligned with the needle prior to injection. Using the previously determined optimal Cas9 amounts for each target, we compared EE and IF obtained by injection into the cell versus the yolk. Our data did not show a significant increase in EE when injecting into the cell (**Figure 2, Supplementary Table 1**). Hence, injection into either the cell or the yolk yields equivalent results.

**Figure 2.**
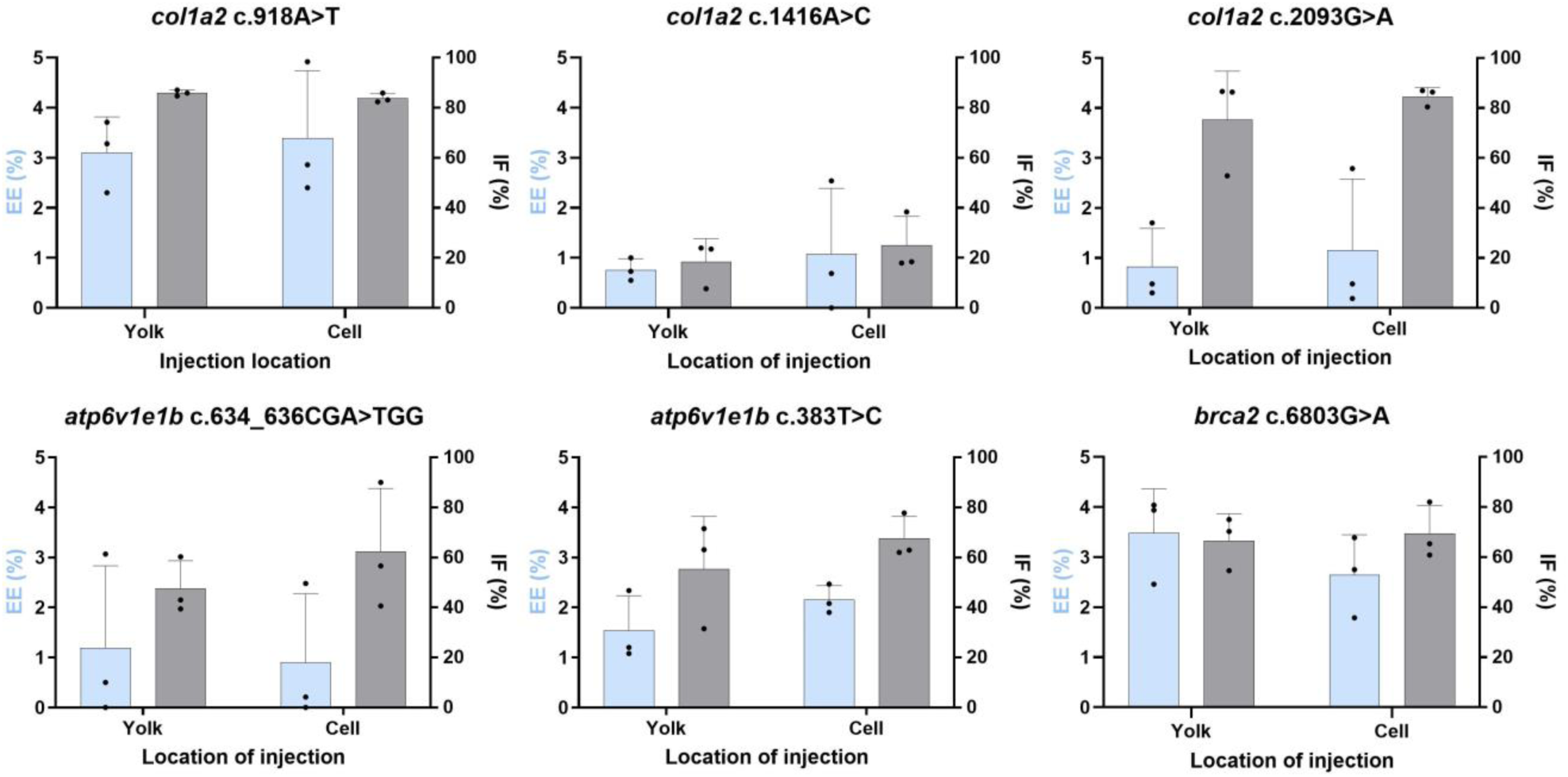
Editing efficiencies (EE - blue) and indel frequencies (IF - grey) obtained by microinjection of Alt-R™ HDR components with 200 pg (*col1a2* c.918A>T), 400 pg (*col1a2* c.1416A>C, *col1a2* c.2093G>A) or 800 pg (*atp6v1e1b* c.634_636CGA>TGG, *atp6v1e1b* c.383T>C, *brca2* c.6803G>A) Cas9 into the cell or the yolk of 1-cell stage zebrafish embryos. Bars and error bars represent the mean + SEM of 3 biologically independent replicates (n=3), where each replicate represents a pool of 20-30 injected embryos. Unpaired t-test.

We used linear regression models to determine the optimal Cas9 amount for achieving the highest EE across all six variants, considering the injection method. For both EE and IF, we evaluated two linear regression models: one including the injection method as a categorical variable and one without it. For EE, analysis of variance (ANOVA) indicated no significant difference between the models (p-value = 0.6044) for injection into the cell versus the less labor-intensive injection in the yolk. Furthermore, post-hoc analysis revealed that higher amounts of Cas9 performed significantly better than lower amounts, with significant differences in EE observed when comparing 100 pg versus 800 pg Cas9 (p-value < 0.001) and 200 pg versus 800 pg Cas9 (p-value < 0.01).

However, for IF, ANOVA showed a significant difference between the models with and without injection method as categorical variable (p-value < 0.05). Injection into the cell resulted, on average, in a modest 7% increase in IF compared to injection into the yolk. Post-hoc analysis also showed significant differences in IF between all Cas9 amounts, with a p-value of 0.01 for the difference between 100 pg and 200 pg, and p-values < 0.001 for all other comparisons. These data suggest that, overall, 800 pg Cas9 induces the most indels (**Figure 2**).

### HDR template modifications result in limited improvements of EE and IF

Modifying the ends of the HDR templates has been reported to increase editing efficiency^19^. One such optimization is the Alt-R™ modification from integrated DNA Technologies. According to IDT, this modification results in higher on-target capabilities compared to unmodified double-stranded DNA with lower non-specific integration elsewhere within the genome (https://www.idtdna.com/pages/support/faqs/what-are-the-modifications-to-the-ends-of-alt-r-hdr-blocks-and-what-is-their-purpose). However, independent studies investigating the effect of these chemical modifications of the HDR template on EE are scarce^19,27^. Although not statistically significant, we observed a trend of lower EE when using unmodified HDR templates (**Figure 3, Supplementary Table 1**) compared to Alt-R™-modified HDR templates (**Figure 3, Supplementary Table 1**). Furthermore, IF appeared to be unaffected by the chemical modifications of the HDR template (**Figure 3, Supplementary Table 1**). In a second experiment silent guide-blocking variants were incorporated into the Alt-R™-modified HDR template. These variants are intended to prevent binding of the crRNA after correct integration of the HDR template and have been reported to increase the EE^28,29^. Surprisingly, we did not observe a significant increase in EE with the incorporation of silent guide-blocking variants. Moreover, introducing a guide-blocking variant significantly decreased the EE for *col1a2* c.918A>T and *atp6v1e1b* c.383T>C (**Figure 3**).

**Figure 3.**
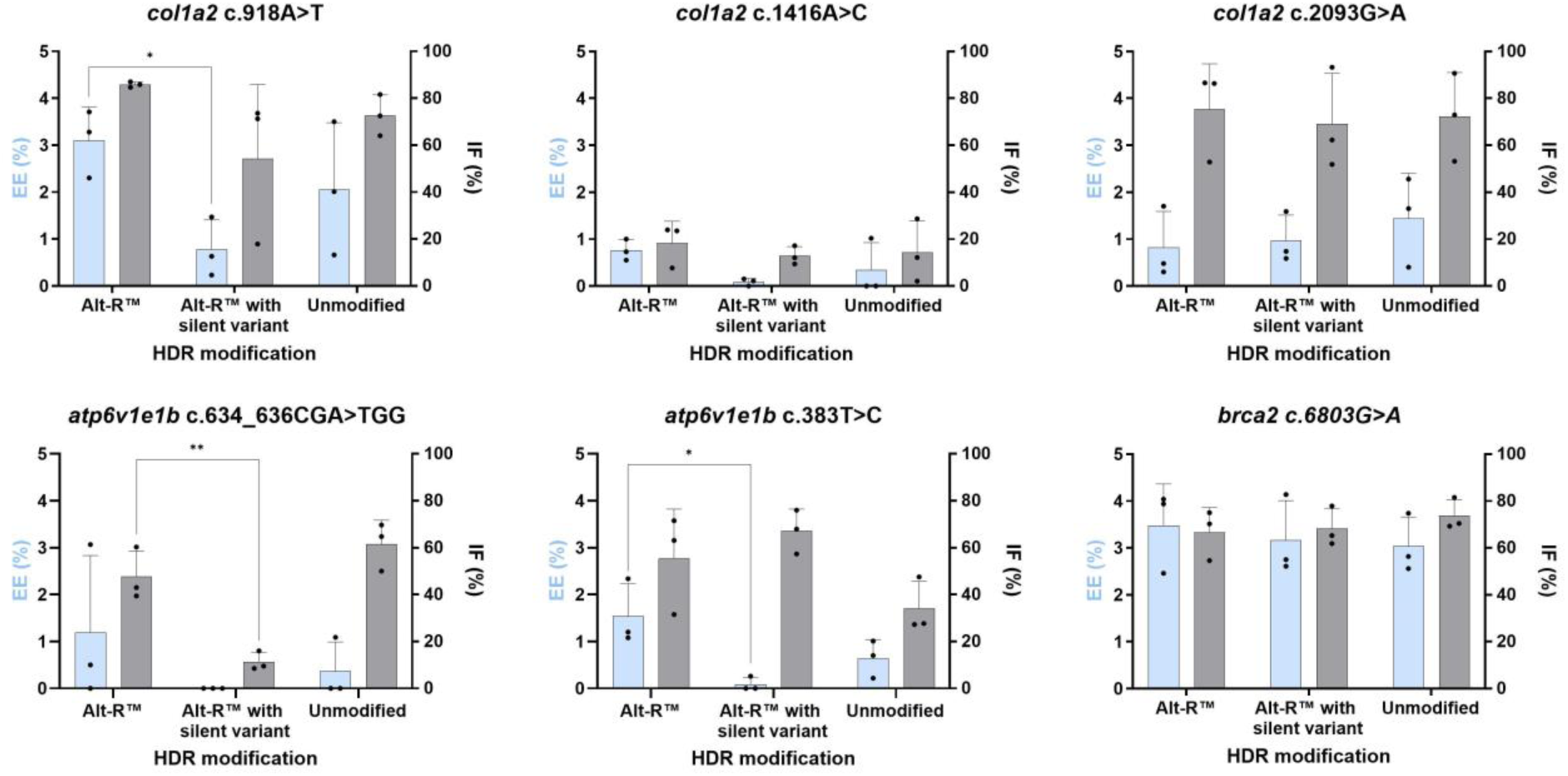
Editing efficiencies (EE - blue) and indel frequencies (IF - grey) obtained by microinjection of Alt-R™ HDR, Alt-R™ HDR with silent guide-blocking variant or unmodified template with 200 pg (*col1a2* c.918A>T), 400 pg (*col1a2* c.1416A>C, *col1a2* c.2093G>A) or 800 pg (*atp6v1e1b* c.634_636CGA>TGG, *atp6v1e1b* c.383T>C, *brca2* c.6803G>A) Cas9 into the yolk of 1-cell stage zebrafish embryos. Bars and error bars represent the mean + SEM of 3 biologically independent replicates (n=3), where each replicate represents a pool of 20-30 injected embryos. Dunnett’s multiple comparisons test: *p < 0.05; **p < 0.01; ***p < 0.001.

Taken together, the optimal injection conditions for the selected targets involve injecting 200 – 800 pg of Cas9 into the yolk along with Alt-R™-modified HDR templates.

### PE outperforms HDR-mediated knock-in in zebrafish

We explored PE as an alternative gene editing method to increase the KI efficiency. To determine if PE achieves higher EE compared to the HDR technique, pegRNAs that introduce the same variants as the HDR templates were designed for all six targets. The PE2:pegRNA RNP complexes were injected in triplicate into the yolk of single-cell zebrafish embryos. We injected 2 nL of the RNP mixture containing 480 ng/µL pegRNA and 1380 ng/µL PE2-His protein. PE was effective for four out of six targets performing equal to (for one target) or better than (for three targets) conventional HDR. A significantly increased EE was observed for *col1a2* c.918A>T (p-value <0.001) and *brca2* c.6803G>A (p-value < 0.05). For *col1a2* c.2093G>A, a trend of increased EE was observed with PE. *Atp6v1e1b* c.383T>C showed similar EE for PE and HDR, while *col1a2* c.1416A>C and *atp6v1e1b* c.634_636CGA>TGG did not show base-pair conversions with an EE higher than 1% using PE. Importantly, indel formation was significantly reduced for all six targets (**Figure 4, Supplementary Table 1**).

**Figure 4.**
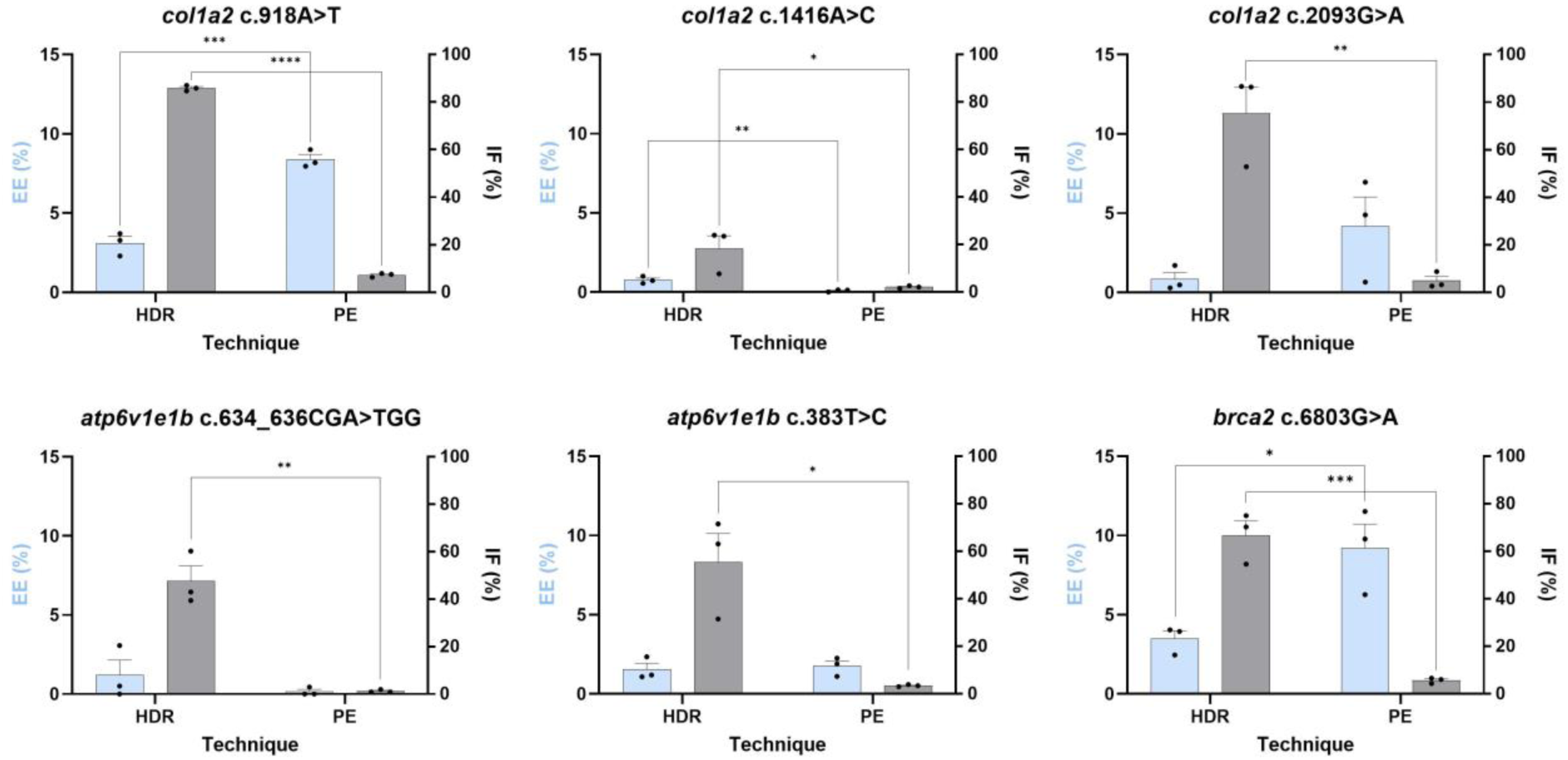
Editing efficiencies (EE - blue) and indel frequencies (IF - grey) obtained by microinjection of either Alt-R™ HDR components with an amount of Cas9 previously determined as the most optimal for that specific target: 200 pg (*col1a2* c.918A>T), 400 pg (*col1a2* c.1416A>C, *col1a2* c.2093G>A) or 800 pg (*atp6v1e1b* c.634_636CGA>TGG, *atp6v1e1b* c.383T>C, *brca2* c.6803G>A) Cas9. EE and IF were compared to injection of PE components into the yolk of 1-cell stage zebrafish embryos. Bars and error bars represent the mean + SEM of 3 biologically independent replicates (n=3), where each replicate represents a pool of 20-30 injected embryos. Unpaired t-test: *p < 0.05; **p < 0.01; ***p < 0.001.

Next, we compared the ability of HDR and PE to introduce variants into the germline using the optimal injection conditions from previous experiments. All fish were grown to adulthood (three months post-fertilization) and the sperm of twenty male zebrafish was screened for correct integration of the desired edit using NGS (**Figure 5, Supplementary Table 2**). Zebrafish with an EE greater than 1% in the germline were labeled as founders. For almost all targets, founders were obtained using PE (5/6), and for most targets (4/6), PE outperformed HDR with up to a 6.5-fold increase in the number of founders detected. Surprisingly, even for *col1a2* c.1416A>C, a founder was obtained using PE despite low EE, earlier determined (**Figure 4**). These founders were able to generate heterozygous offspring.

**Figure 5.**
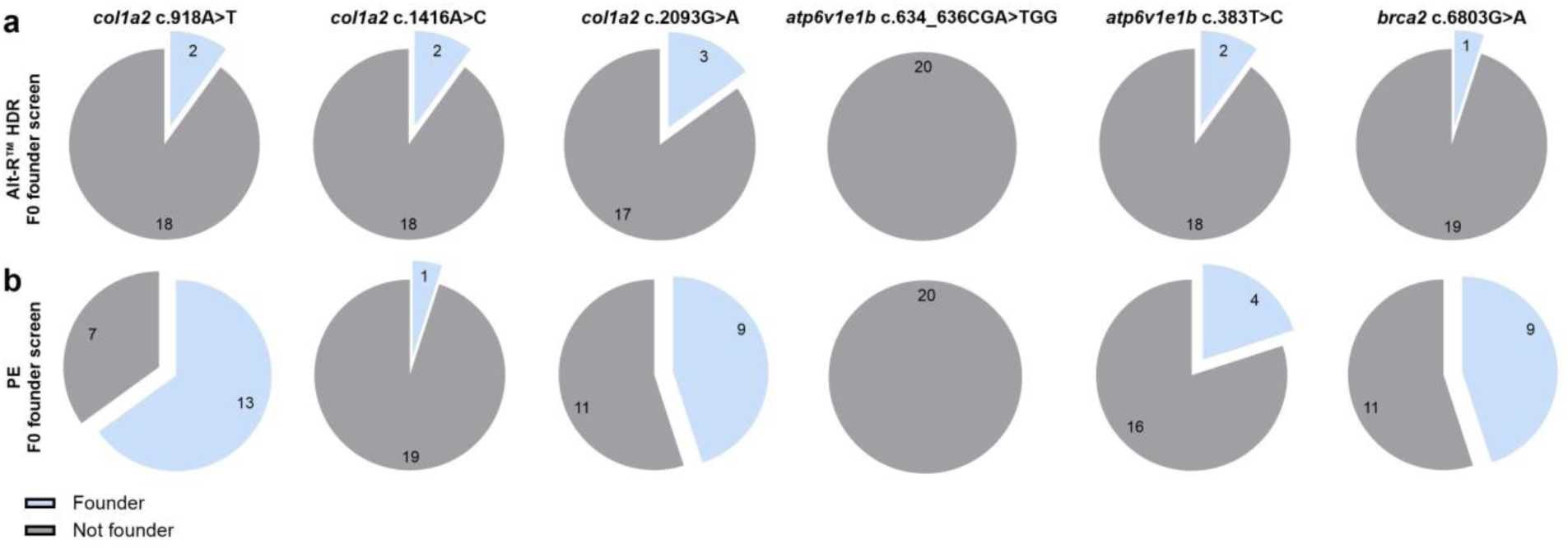
F0 founder screen of 20 potential adult founder zebrafish generated by microinjection of **A)** Alt-R™ HDR components with 200 (*col1a2* c.918A>T), 400 (*col1a2* c.1416A>C, *col1a2* c.2093G>A) or 800 (*atp6v1e1b* c.634_636CGA>TGG, *atp6v1e1b* c.383T>C, *brca2* c.6803G>A) pg Cas9 or **B)** prime editing components into the yolk of 1-cell stage zebrafish embryos.

### PE elicits fewer off-target effects compared to HDR

We used the CRISPOR tool^30^ to predict the three most closely matching genomic off-target sites in the GRCz11 zebrafish reference genome for all guide sequences used (**Supplementary Table 3**). Next, off-target rates were experimentally validated for pools of zebrafish embryos injected with PE or Alt-R™-modified HDR templates using the optimal Cas9 amounts (**Supplementary Table 4**). Among the pools injected using HDR, two gRNAs targeting *col1a2* c.918A>T showed significantly higher IF at off-target site one and three, with an average IF of 8.44% and 6.39% respectively. Importantly, despite high on-target EE for this variant, no off-target effects were observed for these loci using PE. Off-target site two showed no statistical difference of IF comparing HDR and PE. For the three *col1a2* loci and the *brca2* locus, predicted off-target sites were identical for PE and HDR since they utilized the same spacer sequence for the gRNA. For other predicted off-target loci, there was limited indel generation, but at levels comparable to HDR. No reverse transcriptase activity was detected at any off-target loci. For the *atp6v1e1b* loci, no statistics are included since the spacer domains between HDR and PE do not match (**Figure 6, Supplementary Table 4**). Overall, these results support the claim that PE is less susceptible to off-targeting effects compared to HDR.

**Figure 6.**
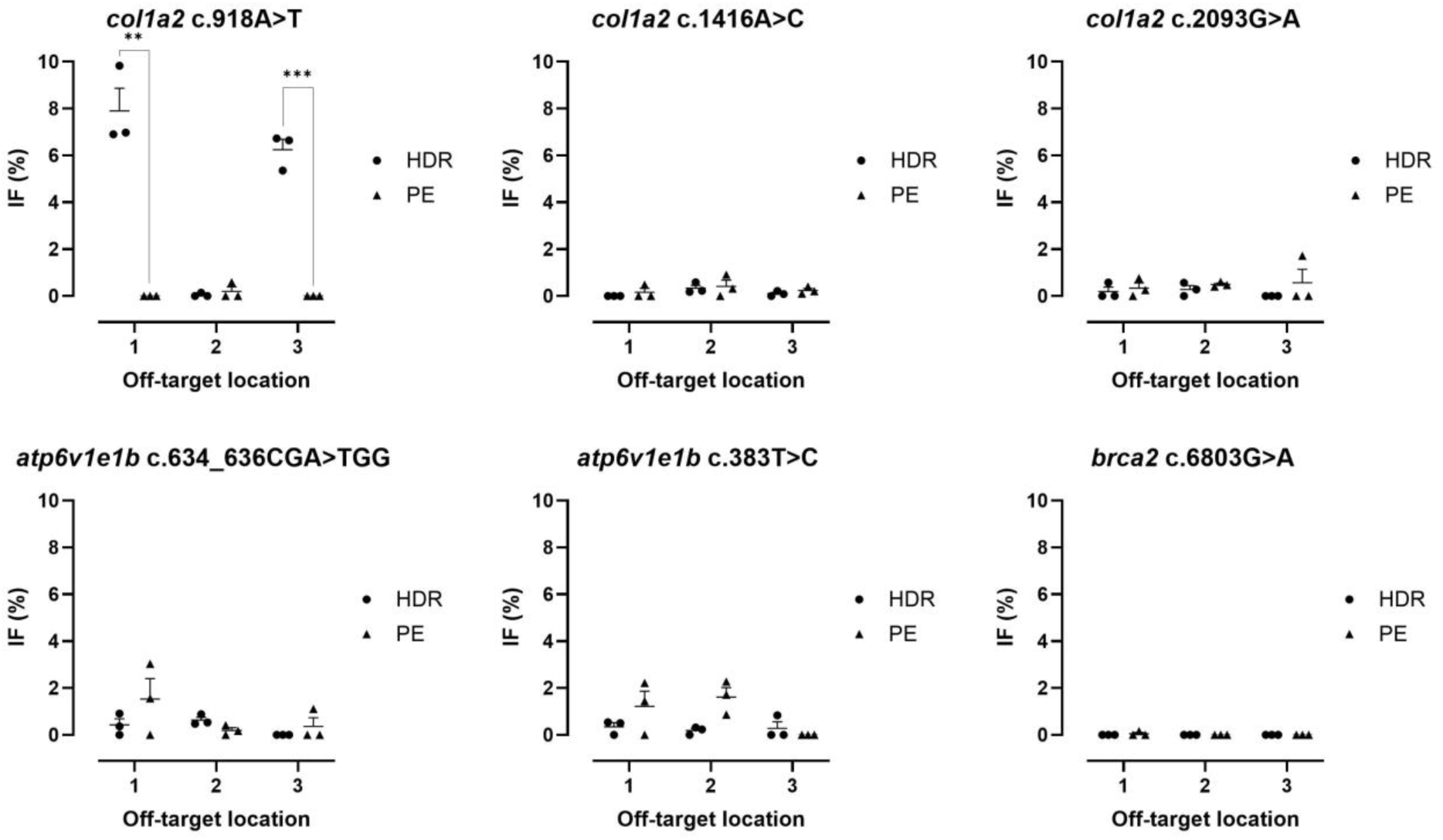
Indel frequency (IF) at top three predicted off-target locations. IF was determined for Alt-R™ HDR components with 200 pg (*col1a2* c.918A>T), 400 pg (*col1a2* c.1416A>C, *col1a2* c.2093G>A) or 800 pg (*atp6v1e1b* c.634_636CGA>TGG, *atp6v1e1b* c.383T>C, *brca2* c.6803G>A) Cas9 or and prime editing components into the yolk of 1-cell stage zebrafish embryos. Bars and error bars represent the mean + SEM of 3 biologically independent replicates (n=3), where each replicate represents a pool of 20-30 injected embryos. Unpaired t-test: **p < 0.01; ***p < 0.001.

## DISCUSSION

Over the past decade, zebrafish have emerged as an attractive model for studying gene functions due to their genetic similarities to humans and advantageous reproductive traits. While traditional KO models are useful, their limitations have driven interest in precise KI models that allow the study of specific human disease-causing variants. However, the efficiency of conventional CRISPR/Cas9 methods, such as HDR, is relatively low. This necessitates not only the optimization of HDR but also the exploration of alternative methods like PE. This study focused on knocking-in six unique variants across three genes associated with Mendelian diseases in humans. The selection of these variants was driven by our goal to create stable mutant KI lines for future research aimed at functionally testing VUS in the *col1a2* and *brca2* genes and assessing the phenotypic effects of known disease-causing variants in the *atp6v1e1b* gene. Our objective was to identify the most efficient method for generating specific variants in the zebrafish genome. For HDR, we selected crRNAs that resulted in the shortest distance between the Cas9 cut-site and the location of the desired edit, as there is an inverse relation between distance and EE^15,31–34^. Unfortunately, using the same spacer domain for PE was not always possible because, for both *atp6v1e1b* targets, the edit is positioned upstream of the cut-site. Therefore, we opted for different spacers that minimized the distance between the edit and the cut-site, while still allowing for PE. Consequently, EE might be affected due to differences in the genetic landscape around the target regions.

We first assessed the impact of different Cas9 amounts on KI EE by HDR, injected as Cas9/gRNA RNP complexes together with Alt-R™-modified HDR templates into single-cell-stage zebrafish embryos. Overall, we observed a trend of significantly higher EE with increased Cas9 amounts (range from 100 to 800 pg). However, establishing a statistically significant optimal Cas9 amount for each target individually proves challenging and labor-intensive. Therefore, it is crucial to minimize variability caused by factors such as droplet size by carefully calibrating the microinjector before each injection experiment.

Despite optimization, EE remains relatively low using HDR in zebrafish. HDR is mainly active during the S and G2 phase of the cell cycle, while NHEJ is active throughout all phases and is a faster repair process compared to HDR^35^. In zebrafish, the first eleven cell divisions occur very rapidly^36^, resulting in short interphases. Due to these fast cell divisions, NHEJ may be favored over HDR^37^. Aksoy *et al* (2019) also showed that slowing down the first cell division by incubation on ice for 15-20 minutes could increase EE, albeit with a lower survival rate, further strengthening this hypothesis^37^. We opted to administer RNPs that, in contrast to Cas9 mRNA, bypass the need for endogenous translation mechanisms, allowing the Cas9 protein to cleave its target more rapidly.

Direct microinjection into the one-cell stage embryo has been reported to yield higher EE^16^ and to be more robust compared to yolk injections^38^. Nevertheless, the short timeframe of the one-cell stage makes intracellular injections technically challenging, as it requires precise positioning and a skilled technician. Injection into the yolk is easier, allowing for rapid and less laborious injections with diffusion of the injected complex into the cell. Comparing efficiencies between component delivery into the cell versus the yolk unexpectedly revealed that intracellular injections do not show a significant increase in EE, while increasing IF by 7% when taking all six targets into account. This suggests that intracellular injection increases cutting efficiency but does not enhance the efficiency of correct integration of the repair template. For KI generation, injection into the yolk may therefore be preferable due to its simplicity and similar efficacy, while being less labor-intensive. In contrast, for KO generation and creating crispant models, which require very efficient indel generation in F0 zebrafish, cell injection is likely to perform slightly better compared to injection into the yolk^39^.

Chemical modifications to HDR templates have the potential to increase the EE (https://www.idtdna.com/pages/support/faqs/what-are-the-modifications-to-the-ends-of-alt-r-hdr-blocks-and-what-is-their-purpose). Additionally, incorporating silent guide-blocking variants into HDR templates is expected to increase EE by preventing re-cutting of the integrated template^29^. In our study, we observed a trend of increased EE with Alt-R™-modified templates compared to non-modified templates. However, the inclusion of a silent guide-blocking variant in the HDR template did not have a significant effect on EE, highlighting the complexity of factors that influence HDR efficiency^29^. The use of silent guide-blocking variants should therefore be approached with caution, since the introduction of an additional (silent) variant may interfere with correct splicing^40–42^. Extra caution should be taken when the incorporation of the silent variant is positioned close to exon-exon junctions, or when new GT or CAG motifs are introduced as these could create new splicing donor or acceptor sites^43^. However, splicing motifs are not always easy to recognize. Exonic splice enhancers are even more difficult to identify and can also result in altered splicing when disturbed^40^. Online tools such as spliceAI^44^ or ESEFinder^41^ can predict the effect of the additional silent variant but are not validated for zebrafish. Therefore, cDNA analysis is advised for the gene of interest in the obtained KI model to detect unexpected splicing.

Notwithstanding improvements to the HDR methodology, KI EE remained low in our study, prompting the exploration of alternative genome editing techniques. Base editing (BE), for example, simplifies the genome editing process by directly converting one base pair to another and has proven successful in different organisms such as yeast, plants, mice, human embryos, and zebrafish^45–48^. However, the scope of BE is restricted to single-nucleotide transitions, which only represent four out of twelve possible single-nucleotide variants. Additionally, multiple cytosines or adenines (depending on the base editor used) within the base editing window are prone to be edited, potentially resulting in unwanted bystander edits. Moreover, this technique does not allow for more complex genomic alterations associated with genetic diseases. Therefore we opted for PE, a recent CRISPR-based genome editing technique that offers greater flexibility, enabling the creation of a wide range of genomic edits, including all possible single-nucleotide variants, DNA insertions, and deletions. In contrast to other methods, PE offers advantages such as precise editing without DSB, reduced off-target effects and increased DNA specificity^20^.

Reports on PE in zebrafish are limited^49–51^, highlighting the need for direct comparisons between HDR and PE for similar targets. Our results confirm the findings of Petri *et al* (2022) that PE2 is functional in zebrafish. They compared the efficiency of KI generation between HDR, using either a low (500 pg per injection) or high (1000 pg per injection) Cas9 amount, and PE in zebrafish for two genes, introducing a deletion, insertion and substitution. Analogous to our results, PE outperformed HDR for the generation of most variants (5/6) and a high dosage of Cas9 scored better in most cases (5/6)^50^. Since only two different spacer domains were explored, and no statistical analysis was provided, their data require validation in additional targets. In our study, PE was effective for five out of the six targets studied and performed equally or better than HDR for germline transmission. For the targets *col1a2* c.1416A>C and *atp6v1e1b* c.634_636CGA>TGG, average EE using PE were 0.093% and 0.015%, respectively (as determined by sequencing of 30 embryos). Nevertheless, by screening twenty adult F0 fish, one founder was obtained for *col1a2* c.1416A>C with 3.37% of the germline containing the desired edit and interestingly, four other potential founders with EE of the germline smaller than 1% were detected. This result indicates that even with very low EE, it is still feasible to obtain founders. In contrast, *atp6v1e1b* c.634_636CGA>TGG showed for all twenty screened fish 0% editing of the germline. A possible explanation for the lack of efficiency of this target could be the short length of the post edit-homology arm. It has been suggested that the length of the post-edit homology arm should have a length of approximately ten nucleotides^50^. For this variant, the longest possible post-edit homology arm using this spacer domain, a PBS of ten nucleotides, and a RTT between twelve and seventeen nucleotides, is four. However, for the *atp6v1e1b* c.383T>C edit, the post-edit homology arm was even shorter, with only three nucleotides and is functional. We hypothesize that the specific genomic landscape around the mutation site has a greater effect on the EE than the pegRNA design, since for both HDR and PE, it was particularly challenging to introduce this variant of interest in the zebrafish genome. Screening a greater number of potential founder fish could increase the chances of finding a founder for this variant, especially since both HDR and PE showed an EE greater than zero when thirty embryos were injected and pooled. Altogether, PE outperformed HDR when comparing the number of zebrafish founders for all targets where PE showed base-pair conversions higher than 1%. This suggests that PE may offer advantages in terms of germline transmission of edited variants, crucial for establishing stable mutant lines. These findings further support the potential of PE as a robust genome editing technique for disease modeling in zebrafish. Further optimization of the PE technique may further improve the EE. One such optimization might be the use of programs that can predict editing rates for all small-sized genetic changes, like PRIDICT (PRIme editing guide preDICTion), which is a neural network optimized in HEK293T and K562 cells. pegRNAs with high (>70) PRIDICT scores showed substantially increased PE EE^52^.

PE significantly reduced unwanted indel formation at both the intended cut-site, and potential off-target locations, compared to HDR. Unlike Petri *et al* (2022), we did not identify off-target integration of the RTT. Conventional HDR raises concerns for unwanted off-target effects on both small and large genomic scale as shown by whole genome sequencing^53^.

In conclusion, our study advances knock-in genome editing in zebrafish. To optimize CRISPR/Cas9-mediated HDR, we assessed the optimal Cas9 amount, injection site, and HDR template modifications to enhance editing efficiency (EE). Additionally, we demonstrated that PE is a superior alternative to HDR, offering precise editing with reduced off-target effects and improved germline transmission rates.

## MATERIALS AND METHODS

### Zebrafish husbandry and maintenance

Zebrafish (*Danio rerio*) were kept in semi-closed recirculating systems (ZebTech, Tecniplast, Italy) at 27 ◦C, pH 7.5, ±550 μS conductivity in a 14/10 light–dark cycle at the Zebrafish Core Facility Ghent (Ghent University). The zebrafish were fed with Zebrafeed (Sparos) and Gemma Micro dry food (Inve) in the morning and Micro Artemia (Ocean Nutrition) in the afternoon. Zebrafish lines were maintained according to standard protocols^54,55^. Zebrafish larvae were raised and placed in an incubator at 28°C, until 5 days-post-fertilization (dpf) with E3 medium. Afterwards, they were transferred to the zebrafish facility and fed on rotifers. The zebrafish AB line was maintained and handled in accordance with the Animal Welfare Legislation, EU Directive 2010/63/EU (European Commission, 2016). The study was approved by the local committee on the Ethics of Animal Experiments (Ghent University Hospital, Ghent, Belgium, permit numbers ECD 22-35K, ECD 22-34, and ECD20-41).

### Generation of gRNAs, ssODN repair templates and pegRNAs

For all six target genes, crRNAs, tracrRNA, ssODN repair templates, and pegRNAs were obtained from IDT. crRNAs were designed using Benchling, prioritizing crRNAs with the shortest distance between the cut-site and edit-site, and to a secondary degree, the highest *in vitro* predicted on-target and off-target scores. The crRNA and tracrRNA were both diluted to a concentration of 200 µM in Duplex buffer (IDT). Next, the two RNA oligos were mixed (1 µL of both) in equimolar concentration to a final duplex concentration of 100 µM, heated for 5 minutes at 95°C and left on the bench top for 10 minutes to cool down to room temperature. Symmetrical ssODN repair templates were designed using the IDT HDR Design Tool with a total length of approximately 120 base pairs (60 nucleotide arms) encompassing the intended edit, with and without Alt-R™ modifications and/or a silent guide-blocking mutations.

pegRNAs were designed using pegFinder^56^ with a PBS length of 10 nucleotides and RTT length between 12 and 17 bp. For the first and last three nucleotides of each pegRNA, phosphorothioated 2’-O-methyl RNA bases were incorporated, which increases the resistance to a variety of ribo-and deoxyribonucleases partially. Standard desalting was chosen as purification method (IDT). Nucleotide composition of all crRNAs, HDR templates and pegRNAs can be found in **Supplementary Table 5**.

### Zebrafish gene editing with HDR

The Alt-R™ S.p. Cas9 Nuclease V3 protein (IDT) was diluted to a 1 µg/µL concentration in a KCl/Hepes buffer (50 mM KCl (Invitrogen) and 11 mM Hepes solution (Sigma) in water and brought to a pH of 7.4). The HDR templates were diluted to a 20 µM concentration in Duplex buffer (IDT). A working solution of 4 µL was made according to volumes described in **Supplementary Table 6**. Absolute weight per injection of the Cas9 nuclease, crRNA:tracrRNA duplex, and ssODN template are provided in **Supplementary Table 7**. Molar ratios per injection (Cas9 nuclease, crRNA:tracrRNA duplex, and ssODN template) are given in **Supplementary Table 8**. Working solutions were incubated for 20 minutes at room temperature and 2µL was loaded in the needle.

Microinjections were performed into the yolk or into the cell of the 1-cell stage AB zebrafish embryos. For every experimental condition, at least fifty embryos per target were injected with 1,4 nL of the RNP mixture and incubated at 28°C. One day post fertilization, at least 30 normally developed embryos were pooled and lysed in 100 µL of 50 mM NaOH at 95°C for 20 minutes. 10 µL of 1 M Tris-HCl was added to the NaOH volume and samples were vortexed and centrifuged at 4000 RPM for 3 minutes. Supernatants containing the DNA was collected and stored at −20°C for subsequent PCR amplification. The remaining injected embryos were kept for monitoring of their further development. Injections were performed in triplicate.

### Zebrafish gene editing with PE

The pET-PE2-His plasmid (IK1822) was a gift from Keith Joung (Addgene plasmid #170103; http://n2t.net/addgene:170103; RRID:Addgene_170103). The PE2-His protein was produced and purified by the VIB protein core (Ghent, Belgium) as previously described, with minor modifications (**Supplementary methods)**^50^. The RNP mixture was made by mixing 1 µL 480 ng/µL pegRNA (in duplex buffer) and 1µL 1380 ng/µL PE2-His protein (in 20 mM Hepes, 150 mM KCl, 10% glycerol at pH 7.5) and incubated for 5 minutes at room temperature. Microinjections were performed into the yolk of the 1-cell stage AB zebrafish embryos. Embryos were each injected with 2 nL of the RNP mixture and incubated at 32°C^50^. Lysates from a pool of 30 embryos were prepared as described above at 1 dpf. Some injected embryos were kept to check for normal development. Injections were performed in triplicate.

### Targeted deep sequencing

The six target site regions were amplified from 2 µL supernatants of the zebrafish embryo lysate with the KAPA2G Fast HotStart PCR Kit (Roche) using the following PCR protocol: (2’ 94°C, 12 x (30’” 94°C, 30” 60*°C, 1’ 72°C), 25 x (40” 94°C, 40” 48°C, 30” 72°C), 10’ 72°C). For each cycle (“*”), the temperature was lowered 1°C. The generated amplicons were then subjected to the Nextera XT library preparation protocol^57^ (Illumina, San Diego, CA) and deep sequenced on a MiSeq instrument (Illumina) using 2×250 bp cycles.

### NGS data analysis

Data analysis was performed using BATCH-GE, a Perl-based bio-informatics tool specifically designed for the assessment of NGS data arising from target-specific genome editing experiments^58^. A window of thirty nucleotides in both directions of the variant site was used. The EE was calculated by dividing the number of reads containing only the intended edit by the total number of mapped reads (pure HDR/PE reads), while IF was calculated by dividing the number of reads with insertions and/or deletions by the total number of mapped reads.

### Screening of founders

HDR- and PE-injected zebrafish embryos for all desired edits were grown until adulthood (3 months post-fertilization). The germlines of twenty potential male founders were screened for the intended edits by extracting and lysing sperm, followed by NGS as described above.

### Off-target detection

Using CRISPOR, the three most closely matching genomic off-target sites in the GRCz11 zebrafish reference genome were identified for all guide sequences^30^. The three highest scoring off-target sites, based on the crRNA, were amplified with genotyping primers and analyzed using NGS as described above. Indel frequencies were calculated using BATCH-GE, using a twenty-nucleotide window around the middle of the off-target crRNA. Data for each potential off-target site are based on injections that were performed in triplicate.

### Statistical analysis

To determine the optimal Cas9 amount that generates the highest EE, a linear regression model was generated in R using the Cas9 amounts applied (100, 200, 400, and 800 pg) and any of the genetic variants studied as categorical variables. The Tukey test was used as a post-hoc test. Optimal Cas9 amounts for integration frequency were also determined with a linear regression model using the Cas9 amounts, the genetic variant, and the injection method (cell or yolk) as categorical variables. Tukey’s honestly significant difference test was used as a post-hoc analysis method. For all other comparisons, GraphPad Prism version 10.1.1 for Windows (GraphPad Software, Boston, Massachusetts USA, www.graphpad.com) was used. For the comparisons between different amounts of Cas9, as well as different HDR template modifications, we employed one-way ANOVA followed by Tukey’s Multiple comparisons test for cas9 amounts and Dunnett’s multiple comparison tests for comparison between Alt-R™-modified HDR templates, non-modified HDR templates and templates with incorporated silent guide-blocking variants. For the comparisons between cell versus yolk injection and PE versus HDR, unpaired Student’s t-tests were utilized. The null hypotheses were rejected at the P < 0.05 level.

## ACKNOWLEDGMENTS

The authors thank the Zebrafish Facility Ghent Core at Ghent University, and particularly Karen Vermeulen for the diligent care for the zebrafish. We would also like to thank the VIB protein core for the generation of the prime editor and professor Joung for the donation of the pET-PE2-His plasmid (Addgene plasmid # IK1822).

## SUPPLEMENTARY MATERIAL

Supplementary table 1

Supplementary table 2

Supplementary table 3

Supplementary table 4

Supplementary table 5

Supplementary table 6

Supplementary table 7

Supplementary table 8

