## Supplementary Methods for "Prime editing outperforms homology-directed repair as a tool for CRISPR-mediated variant knock-in in zebrafish"

***Supplementary methods: Generation of the PE***

The C-terminal His-tagged PE2 protein was overexpressed in Rosetta 2 (DE3) competent cells and transformed via heath shock using manufacturers protocols. A single colony was picked and grown overnight at 37°C in Luria Bertani (LB) medium with 50 µg/mL kanamycin and 100 µg/mL chloramphenicol. This culture was subsequently used to inoculate 12 L of LB medium with 50 µg/mL kanamycin at 37°C until the optical density (OD600) of the culture had reached 0.65. The expression was induced with 1 mM isopropyl ß-d-1-thiogalactopyranoside (IPTG) at 20°C for 5 hours and overnight at 28°C. Cells were then harvested and frozen at -20°C. After thawing, cells were resuspended at 3 mL/g in lysis buffer 50 mM Tris(Hydroxymethyl)aminomethane pH 8, 1 M NaCl, 1 mM Dithiothreitol (DTT), 1 tablet/200 mL cOmplete Protease Inhibitor Cocktail EDTA-free (Roche). The cytoplasmic fraction was prepared by sonication of the cells followed by centrifugation at 18,000 g for 30 min. All steps were conducted at 4°C. The lysate was treated with 5% polyethyleneimine (Serva) to precipitate host DNA. The supernatant was isolated for purification by centrifugation at 18,000 g for 30 min. The supernatant was then treated with 70% ammonium sulphate to precipitate the protein fraction. The protein fraction was resuspended is loading buffer 2 x phosphate buffer saline (PBS), 20 mM imidazole, 1 mM DTT and applied to a 6 mL Ni-Sepharose 6 FF column (GE Healthcare). After equilibration with loading buffer during 5 column volumes, there was an intermediate wash step with 6 x PBS during 5 column volumes followed again with loading buffer during 10 column volumes with finally an elution in 2 x PBS, 250 mM imidazole, 10% glycerol and 1 mM DTT. The elution fraction was diluted with 50 mM Hepes pH 7.5, 10% glycerol to a conductivity of 11 mS/cm and loaded on a 12 mL Capto S column (GE Healthcare) to remove contaminants. After equilibration with 50 mM Hepes pH 7.5, 10 % glycerol, 100 mM NaCl; the protein of interest was eluted by a linear gradient over 15 column volumes of NaCl from 100 mM to 300 mM followed by a regeneration of the column with 1 M NaCl. Finally, the recombinant protein was dialyzed to with 20 mM Hepes pH 7.5, 10% glycerol, 150 mM KCl. The obtained purified protein was analyzed by SDS-PAGE and the concentration was determined using via Little Lunatic (UV280 nm) at 1.38 mg/mL. Solution was filter sterilized before the concentration was measured and stored at -80°C.
